# Role of diffusion and reaction of the constituents in spreading of histone modification marks

**DOI:** 10.1101/2023.12.13.571420

**Authors:** Vinoth Manivannan, Mandar M. Inamdar, Ranjith Padinhateeri

## Abstract

Cells switch genes ON or OFF by altering the state of chromatin via histone modifications at specific regulatory locations along the chromatin polymer. These gene regulation processes are carried out by a network of reactions in which the histone marks spread to neighboring regions with the help of enzymes. In the literature, this spreading has been studied as a purely kinetic, non-diffusive process considering the interactions between neighboring nucleosomes. In this work, we go beyond this framework and study the spreading of modifications using a reaction-diffusion (RD) model accounting for the diffusion of the constituents. We quantitatively segregate the modification profiles generated from kinetic and RD models. The diffusion and degradation of enzymes set a natural length scale for limiting the domain size of modification spreading, and the resulting enzyme limitation is inherent in our model. We also demonstrate the emergence of confined modification domains without the explicit requirement of a nucleation site. We explore polymer compaction effects on spreading and show that single-cell domains may differ from averaged profiles. We find that the modification profiles from our model are comparable with existing H3K9me3 data of *S. pombe*.

## I. INTRODUCTION

Even though all cells in a multi-cellular organism have the same genetic code (DNA sequence), the different cell types in the organism, for example, neuronal cells and skin cells, behave very differently. This difference in function is achieved by encoding extra layers of information in the form of an epigenetic code [1, 2]. The significant players in encoding this epigenetic code are nucleosomes — 147 bp of DNA wrapped around an octamer of histone proteins — positioned throughout the genome [3–5]. The functional genome is a chromatin polymer with nucleosome as its fundamental unit and other proteins helping in the assembly and disassembly of nucleosomes, alterations in chemical states of the histones, and 3D organization of DNA [6–11].

While the positions of nucleosomes along the DNA itself carry some epigenetic information, more information is encoded in the form of chemical modifications on the histones [2, 12–15]. For example, the 9^th^ amino acid lysine of histone H3 gets tri-methylated (H3K9me3) or acetylated (H3k9ac), and the presence of these marks along the chromatin polymer contour decides whether the local chromatin is inactive or active. Research in the last two decades has given us a basic understanding of how different chromatin marks affect the state of the local chromatin, including the 3D folding, accessibility of certain regions, and physiological state of the cell [16–22]. For instance, it is known that abnormal histone modifications can lead to various diseases, such as bone and brain cancer [23, 24]. While we understand the steady-state profiles of the histone modifications from CHIP-seq experiments [25, 26], very little is known about the kinetics of the formation of these profiles [27–29]. Therefore, it is important to understand these underlying mechanisms of histone modification pattern establishment.

Following DNA replication, the *de novo* assembly of chromatin states involves nucleation of histone modifications at specific target sites [30–33]. Experimental evidence shows that around the nucleation site, the modifications are able to spread to the neighboring regions and establish a stable pattern of modifications [32, 34, 35]. Specifically, H3K9me3 has served as a paradigm to study the spreading of histone modifications. Hathaway *et al*. studied the spreading of H3K9me3 from Oct4 locus [36], and showed that the modification starts near a nucleation site and is confined to a limited region along the chromatin contour. In *Schizosaccharomyces pombe*, the mating type locus is repressed by the spreading of H3K9me3 [37, 38]. In this case, the histone methyltransferase complex (“reader/writer”) Clr4/Suv39h recognizes H3K9me3 and spreads the mark throughout the domain via RNAi mechanism [39, 40]. Recently there have been experiments that probe the heterochromatin assembly and the signaling pathways that govern it [41–43].

Many theoretical models were proposed to study the positive feedback of reader-writer enzymes in establishing chromatin domains [44–51]. Dodd *et al*. provided a model that investigates the heritability and bistability of acetylations and methylations at various locations along the genome [47]. Another model was proposed by Hodges *et al*. [52], where the spreading starts from the nucleation point and proceeds in a set of kinetic events, leading to a modification pattern that decays away from the nucleation point. Following these, many groups have explored different sets of kinetic events and non-equilibrium rules, studying the spreading and formation of histone modification patterns [53, 54]. The spreading along the linear dimension is the most natural event, given the beads on a string picture of chromatin. However, 3D spatial proximity between far-away nucleosomes could also be important, and recent papers have incorporated this into their models to explain the spreading [55–61].

If modifications can spread to all proximal nucleosomes in 3D, it is not clear exactly how a modification limits itself into domains of finite size. Some hypotheses have been proposed to address this question; they include having boundary elements that slow down the spreading [31, 54, 57, 62], looping mechanisms that relax chromatin quicker [55–60], the possibility of multiple intermediate states [63], the presence of large gaps between sliding nucleosomes [64–68], and enzyme limitation [61, 69].

Even though several papers study the spreading of histone modifications computationally, no model so far has explicitly considered the diffusion of the enzymes and RNA that are crucial in the spreading process. Existing models assume that the process is reaction-limited. However, two essential points concern this assumption: (i) Recent experiments have shown that the viscosity inside the nucleoplasm can be very high— orders of magnitude higher than water [70]. This implies that the diffusion can be very slow and can influence the effective time scale of the reactions [71]. (ii) Irrespective of the diffusion time scale, kinetic parameters such as decay of the enzymes/RNA-complexes can limit the spreading process.

In this paper, we examine the interplay between diffusion of enzymes/complexes and reaction-kinetic events in deciding the spreading of histone modifications. We propose a reaction-diffusion framework to study the establishment of modification domains following the nucleation events. We explicitly consider the diffusing factors and the production and decay of the constituents involved in the spreading process. We find that a purely kinetic model may not always explain the modification profile of an entire reaction-diffusion system. We also examine the effect of polymer folding and far-away contacts in our model.

## II. MODEL

### A. Reaction-Diffusion Model

We study the spreading of the histone modifications in a system consisting of a few kilobase (kb) of chromatin (*N*_*n*_ nucleosomes), enzymes that read/write histone modifications, and RNA. We use a particle-based reaction-diffusion model to understand the spreading of the histone modifications from a nucleation point. We choose a circle of radius *r* as our simulation space with *N* particles in it. The particles can be any one of the three types: Enzyme (*E*), RNA (*R*) or Enzyme-RNA complex (*C*). The diffusion of the particles in the system is simulated using Langevin dynamics (see eq. 1). Given the position of *i*^*th*^ particle at the *n*^*th*^ timestep (**r**_**i**_(*n*)), we update it for the next time step as follows:

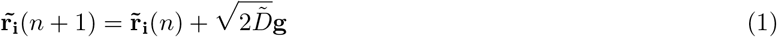

where 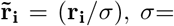 diameter of a particle, 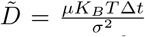 is the dimensionless diffusion constant, *µ*= mobility of particles, *K*_*B*_= Boltzmann constant, *T* = Temperature, and *g* is a Gaussian random number with mean 0 and variance 1. We consider only the thermal kicks obeying the fluctuation-dissipation theorem. We assume that all particles have the same size and diffusion constant for simplicity,.

Experimental evidence so far suggests that most of the DNA folding proteins (HP1, PRC, CTCF) bind at locations with specific histone modifications [72–74]. This implies that the spread of histone modifications, to some extent, would precede the 3D folding of chromatin. Hence, it is crucial to understand the kinetics of 1D spreading. Therefore, we consider chromatin as a 1D polymer in the first part of the work, assuming it is linear in the scale of a few nucleosomes and the most critical spreading events occur between the neighboring nucleosomes. In a later section (see section III E), we present results considering chromatin as a 2D polymer, examining the effect of the folding/compaction. In the 1D polymer picture, each nucleosome is a lattice point along the x-axis with its origin at the center of the circle (see Fig. 1). The lattice spacing between neighboring nucleosomes is unity (*σ* units). For simplicity, each nucleosome can be either in Unmodified (U) or Modified (M) state. The nucleosome at the origin is considered the nucleation point (NP) and maintained at M state throughout the simulations. In fission yeast, there is evidence that the Clr4/Suv38h complex recognizes the existing modification in a nucleosome and propagates it to the neighboring nucleosomes [75]. The localization of these complexes is observed to be through RNAi mechanism [18, 38–40, 75]. RNAi mechanism involves cleaving of mRNAs generated to form siRNAs, and then siRNAs become an integral part of these enzyme complexes. This is used by the cell as a localization mechanism for heterochromatic silencing [37].

**FIG. 1:**
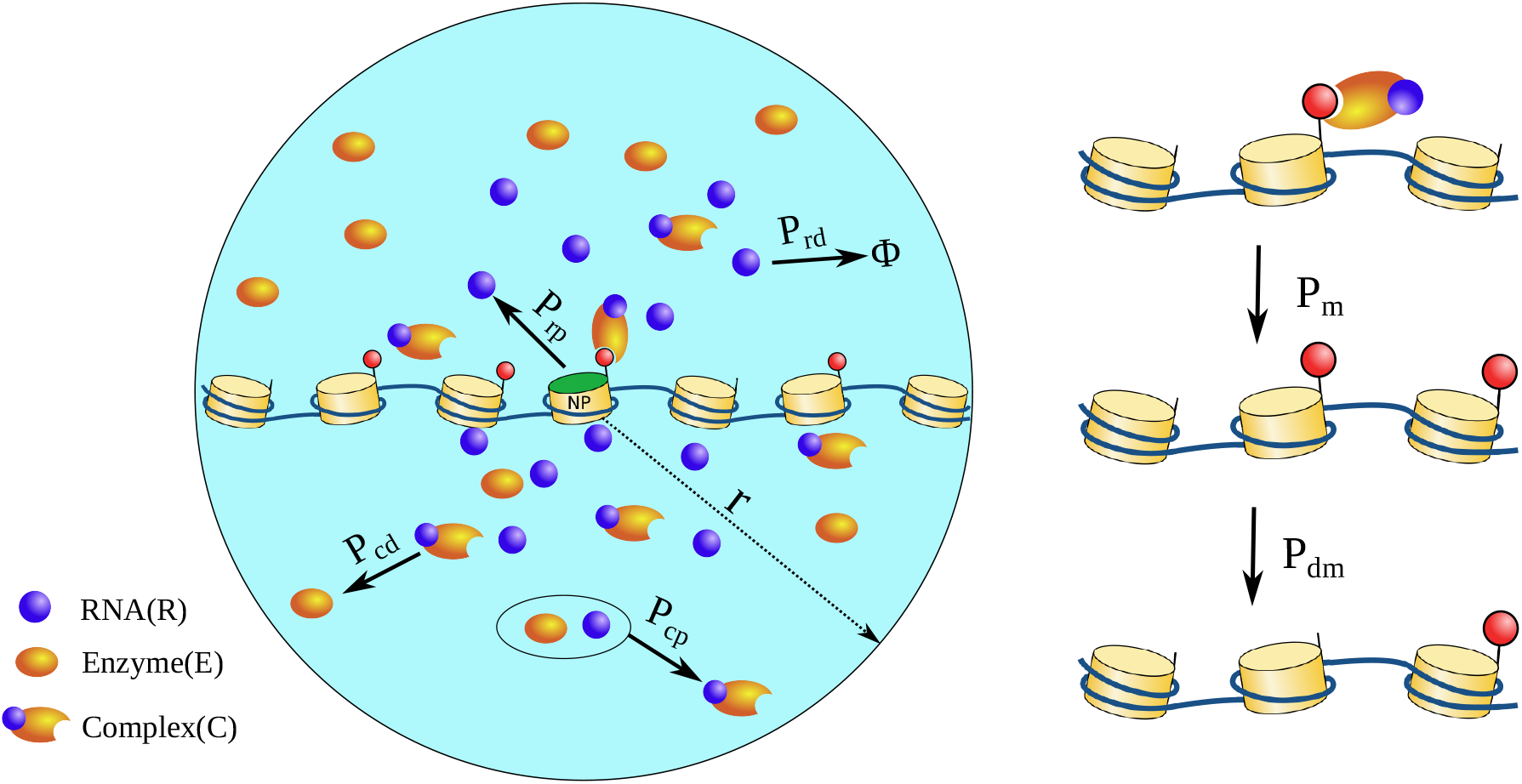
Schematic of the reaction-diffusion model in which each lattice site represents a nucleosome. Particles *R, E*, and *C* diffuse in the simulation space. *R* is produced only at the nucleation point (NP) with probability *P*_*rp*_. It can decay with probability *P*_*rd*_. When *R* and *E* come in proximity, they form *C* with probability *P*_*cp*_. *P*_*cd*_ controls the decay of *C* back to *E* (see eqs. 2 to 4). The *C* particles spread the modification to the nearest neighbor with probability *P*_*m*_ (see right-hand side). Each modified nucleosome can become unmodified with probability *P*_*dm*_.

To simulate the essence of the mechanism described above, we consider a simple scenario of RNA-like particles generating from the origin, which on collision with enzyme particles form enzyme-RNA complexes. The reactions involved in the model are:

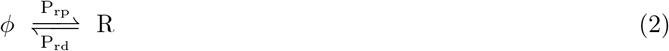

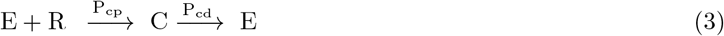

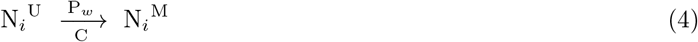

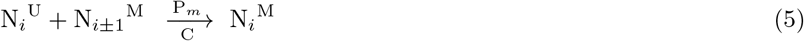

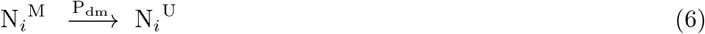

RNA (*R*) is being produced only at the origin, with the probability *P*_*rp*_, and it can decay to *φ* with the probability *P*_*rd*_ anywhere in space as it diffuses. Enzyme particles (*E*) are uniformly distributed in space at the start of the simulation. When *E* comes in contact with *R*, they form complex (*C*) particles with the probability *P*_*cp*_. Enzyme-RNA complex (*C*) particles can methylate a nucleosome with a probability *P*_*w*_. Also if one of the nucleosome’s neighbors (*N*_*i±*1_) is methylated, the probability of it getting methylated will be boosted by *P*_*m*_. Therefore, the effective probability of methylation will be, 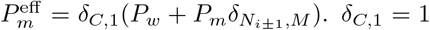 if *C* complex is present in the vicinity of a nucleosome, otherwise 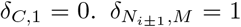 if either of neighboring nucleosomes is methylated, else 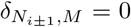. The value of *P*_*w*_ is non-zero in only the results presented in section III D.

### B. Model with only kinetic events

To compare our results of the reaction-diffusion model with existing models in the literature, we also simulate the dynamics of the spreading of histone modifications as a purely kinetic process, similar to the work done by Hodges *et al*. [52]. The nucleosomes are considered as a string of lattices (*j* ∈ [ *−* 30, 30]), which can stay either in a modified (M) or unmodified state (U). The lattice position “0” is assumed as the nucleation point, which is always maintained at the modified state. The kinetic events considered are: (i) Spreading: *k*^+^ represents the rate of addition of the marks to the lattices’ neighbors that are already modified. (ii) Turn-over: *k*^*−*^ represents the rate of removal of marks from any lattice. Given a set of parameters, the stochastic evolution of these states of the nucleosomes is simulated by a standard Gillespie Monte-Carlo algorithm [76–79].

## III. RESULTS

Our aim is to study the role of diffusion and the reaction of enzymes and other necessary constituents in the spreading of histone modifications along the chromatin polymer chain. To do this, we consider a chromatin region having *N*_*n*_ nucleosomes with a nucleation site, mRNA production, and reaction-diffusion mimicking RNAi mechanism along with enzyme diffusion and reactions. We modeled the chromatin region as a string of a one-dimensional lattice of 61 nucleosomes (index, *I* ∈ [ *−* 30, 30]), where they can exist in the modified (e.g. methylated) or unmodified (e.g. unmethylated) state (see Fig. 1). Since our interest is to study methylation spreading, for simplicity, we only consider methylated or unmethylated states. The simulation box contains *N*_*e*_ number of enzyme particles (E) that are uniformly distributed at the start of the simulation. During the simulation, RNA-like particles (*R*) are produced from a specific location (i=0, also the nucleation point NP) on the chromatin (see Fig. 1). The particles (*E* and *R*) diffuse in the simulation box, obeying Langevin dynamics. When they come in proximity, they react with a probability to form enzyme-RNA complex (*C*) particles (see Fig. 1 and section IIA). When *C* is near an unmodified nucleosome and if either of its neighbors is in the modified state, the nucleosome can get modified with a probability *P*_*m*_ (see eq. 5). Any modified nucleosome can become unmodified with a probability *P*_*dm*_ (see eq. 6). During the simulation, we record the modification state of each nucleosome (methylated or unmethylated) and analyse it statistically.

### A. Enzyme-RNA complex and RNA reaction parameters influence the methylation profile

Using an ensemble of snapshots from our simulation—modification of all nucleosomes—at the steady state, we compute the probability that the nucleosome at a given lattice site is methylated (*P* (*M*)). This site-dependent methylation probability would be referred to as the methylation profile in this manuscript. In this section, we vary the enzyme and RNA reaction parameters and investigate how the resulting methylation profile varies. In Fig. 2A, we show the methylation profile by varying only the RNA decay probability *P*_*rd*_ (see eq. 2) while keeping the remaining parameters constant. We see that the increase in *P*_*rd*_ from 10^*−*5^ to 10^*−*3^ significantly affects the spreading of the methyl marks, as inferred from the wider methylation profile. The spreading of the methyl marks is also quantified in the inset by plotting the standard deviation (*S*_*x*_) of the methylation profile for different sets of parameters,

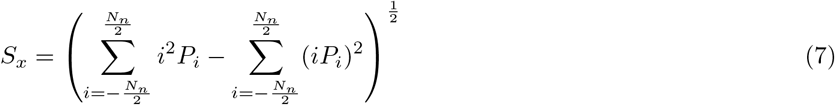

where *i* = the lattice position of the nucleosome, and *P*_*i*_ corresponds to the probability of modification (*P* (*M*)) of the lattice *i*. Fig. 2B shows that at higher *P*_*dm*_ values, change in *P*_*rd*_ does not affect the spread of the methylation. Similarly, in Fig. 2C, we plotted the methylation profile by varying the complex decay probability*P*_*cd*_ (see eq. 4). As *P*_*cd*_ decreases, the spreading increases in the case of *P*_*dm*_ = 10^*−*3^. In contrast, it remains unchanged when *P*_*dm*_ = 10^*−*2^ (see Fig. S1). This implies that *P*_*dm*_ = 10^*−*2^ dominates the effects of *P*_*rd*_ and *P*_*cd*_. Meanwhile, when *P*_*dm*_ = 10^*−*3^, we can see a competition between these values to spread the methyl marks. A phase diagram of the time evolution of modifications in the nucleosomes and the resulting profile is shown in Fig. S2 and Fig. S3, as we vary *P*_*rd*_ and *P*_*cd*_.

**FIG. 2:**
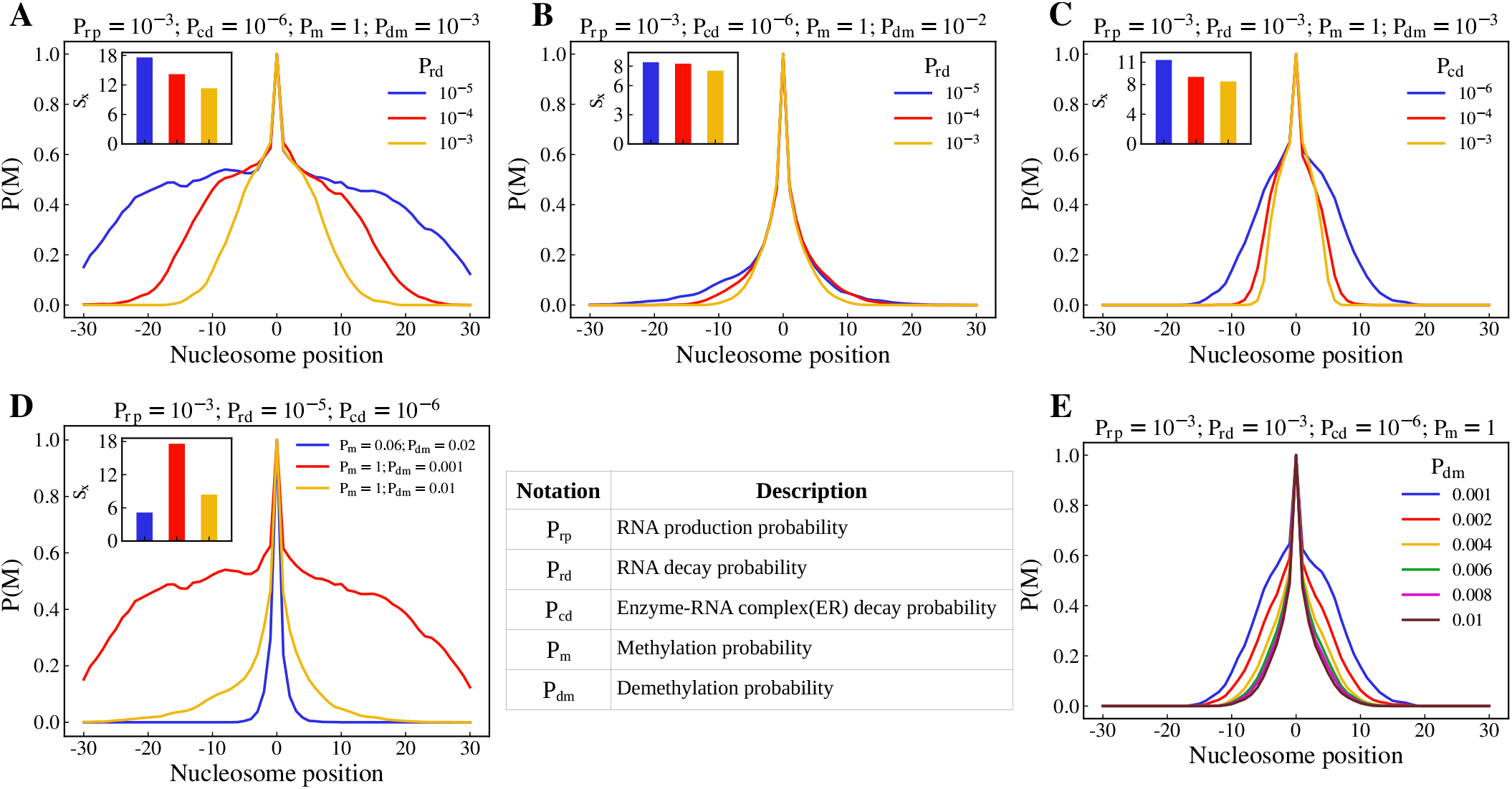
Effect of kinetic parameters on the methylation profile. Methylation profile compared by varying *P*_*rd*_ when *P*_*rp*_ = 10^*−*3^, *P*_*cd*_ = 10^*−*6^, *P*_*m*_ = 1 (A) *P*_*dm*_ = 10^*−*3^. (B) *P*_*dm*_ = 10^*−*2^. (C) Methylation profiles compared by varying *P*_*cd*_ when *P*_*rp*_ = 10^*−*3^, *P*_*rd*_ = 10^*−*3^, *P*_*m*_ = 1, *P*_*dm*_ = 10^*−*3^ (D) Methylation profiles by varying *P*_*m*_ and *P*_*dm*_. (E) Methylation profiles while varying *P*_*dm*_ from 10^*−*3^ to 10^*−*2^, where all the other parameters are constant

The enzyme-mediated modification reactions of the nucleosomes occur stochastically at certain rates. The reactions have two steps. In the first step, the reactants diffuse and reach near the nucleosomes. Even after reaching the proximity of nucleosomes, the reactions are not immediate; it has a stochastic nature and hence occur with a methylation probability*P*_*m*_ (see eq. 5) in our simulations. The demethylation reaction is accounted for by a probability*P*_*dm*_. In Fig. 2D, we compare the methylation profile from three sets of simulations that have the same reaction parameters for R and C (*P*_*rp*_ and *P*_*rd*_, *P*_*cp*_ and *P*_*cd*_), but different *P*_*m*_ and *P*_*dm*_ (See subfigures’ title for the exact values of the parameters). We see that increasing *P*_*m*_ from 0.06 to 1 results in a slightly wider methylation profile, whereas decreasing *P*_*dm*_ from 10^*−*2^ to 10^*−*3^ results in the spread of the methyl marks to the extremities of the lattice. We vary only *P*_*dm*_ in Fig. 2E while keeping all the other parameters constant. The methylation profile gets narrower as we increase *P*_*dm*_.

We also show the effects of the reaction parameters for the production and decay of *R* and *C* (*P*_*rp*_, *P*_*rd*_, *P*_*cp*_, *P*_*cd*_) in the spatial distribution of *R* and *C*. To quantify the spatial distribution of RNA, we compute the number density of RNA (see Fig. S4) at radius *r*_*i*_ as,

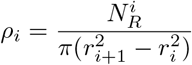

where, 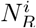 is the number of RNA in the *i*^*th*^ shell and *r*_*i*_ is the radius of the *i*^*th*^ circle.

Fig. 3A shows the number density of RNA in the *i*^*th*^ shell, calculated when *P*_*rp*_ = 10^*−*3^. The RNA profile gets narrower as we increase *P*_*rd*_. Also, we can notice that there is no difference in the RNA profile by changing the value of *P*_*rp*_ to 10^*−*4^. Therefore, we have kept the value of *P*_*rp*_ = 10^*−*3^ throughout this paper. The width of the profile represents the length scale (*l*_*r*_) beyond which RNA cannot be found and can be estimated as 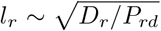, where *D*_*r*_ is the diffusion coefficient of the RNA particles. When E and R particles come in proximity, the enzyme-RNA complex (*C*) is formed with a probability*P*_*cp*_ (see eq. 3). The mean number of *C* particles proximal to the nucleosomes is determined by the RNA length scale (*l*_*r*_) and the probabilities *P*_*cp*_ and *P*_*cd*_. Since the meeting of the E and R is less likely, we have assumed the formation of *C* to be diffusion-limited, by keeping the *P*_*cp*_ = 1 in the results we present. Fig. 3B shows the mean number of complex particles proximal to the nucleosomes for different *P*_*cd*_ values, when *P*_*rd*_ = 10^*−*3^. At higher *P*_*cd*_, the complex particles are more localized to the nucleation point. The same quantity for simulations with *P*_*rd*_ = 10^*−*4^ is plotted in Fig. 3C. We can see that profiles of the complex particles in Fig. 3C are wider than the corresponding profiles in Fig. 3B, which is due to the higher *P*_*rd*_ value of the latter.

**FIG. 3:**
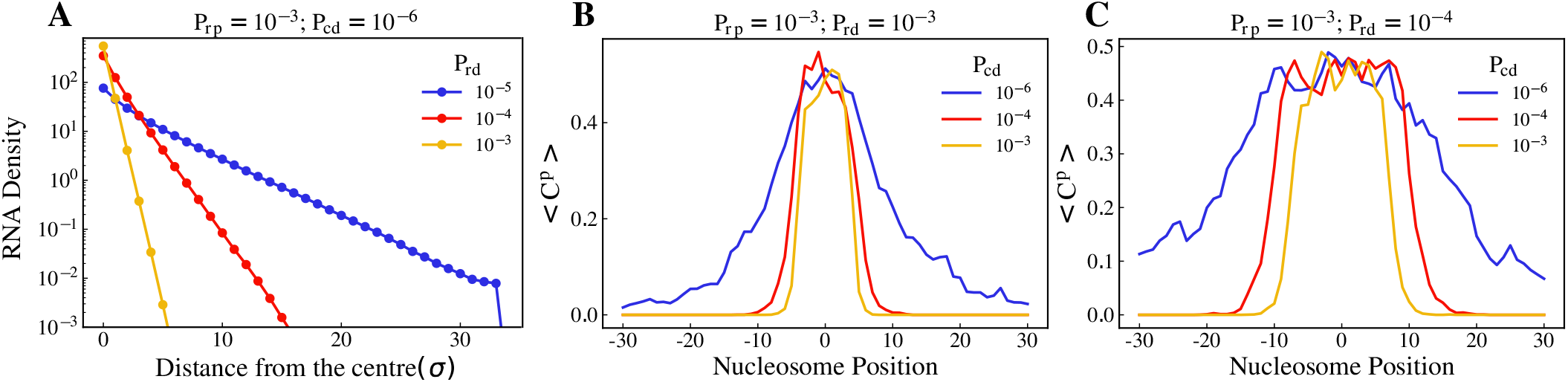
Spatial distribution of *R* and *C* at the steady state. (A) Number density of R present in the *i*^*th*^ concentric shell (*ρ*_*i*_) for different values of *P*_*rd*_, *i* = 0 refers to the center of the simulation space (see text) when *P*_*rp*_= 10^*−*3^. (B) Average number of C particles proximal to the nucleosome position (*< C*^*p*^ *>*) is plotted against nucleosome position for different *P*_*cd*_ values when *P*_*rd*_= 10^*−*3^. (C) when *P*_*rd*_ = 10^*−*4^.

### B. Enzyme limitation influences the methylation profile

Conventional kinetic models assume enzyme supply to be infinite, and the reactions go on at a constant rate forever. Typically, in cells, the amount of protein present is finite. Accounting for that is referred to as enzyme limitation. Since our model explicitly considers a finite number of diffusing protein particles, enzyme limitation is naturally incorporated into our model. We investigated enzyme limitation in two ways: (i) varying the radius of the system (area of the simulation region) fixing the number of enzyme particles (*E* and *C*) constant, effectively varying the concentration of the enzymes, and (ii) changing the total number of the constituents (proteins and RNA), fixing the area of the simulation region constant.

For case (i), we simulated a 41-nucleosome lattice with 500 enzyme particles (*N*_*e*_) for a given set of reaction parameters and varied only the radius of the simulation region, as mentioned above. In Fig. 4A, we show the methylation profiles for the system radii ranging from 25*σ* to 60*σ*. The spread of the methyl marks is dependent on the concentration of the particles, which is inversely related to the radius of the simulation region. In Fig. 4B, the standard deviation *S*_*x*_ (see eq. 7) is plotted against the concentration of enzyme in the system (concentration = *N*_*e*_*/πr*^2^), where *N*_*e*_ is the number of enzyme particles in the system and *r* is the radius of the system. It increases with increasing enzyme concentration (i.e., reduction of radii). The resulting concentration profile for the complex is shown in Fig. S5. The spreading is close to zero at any concentration below 0.1. This elucidates the importance of the local concentration of the enzymes in the spreading. For case (ii), we compared the methylation profiles for the number of enzyme particles (*N*_*e*_) as 500 and 1000, which is shown in Fig. 4C. Despite having the same reaction parameters, the *N*_*e*_ = 1000 particle system has a wider methylation profile than the *N*_*e*_ = 500 particle system.

**FIG. 4:**
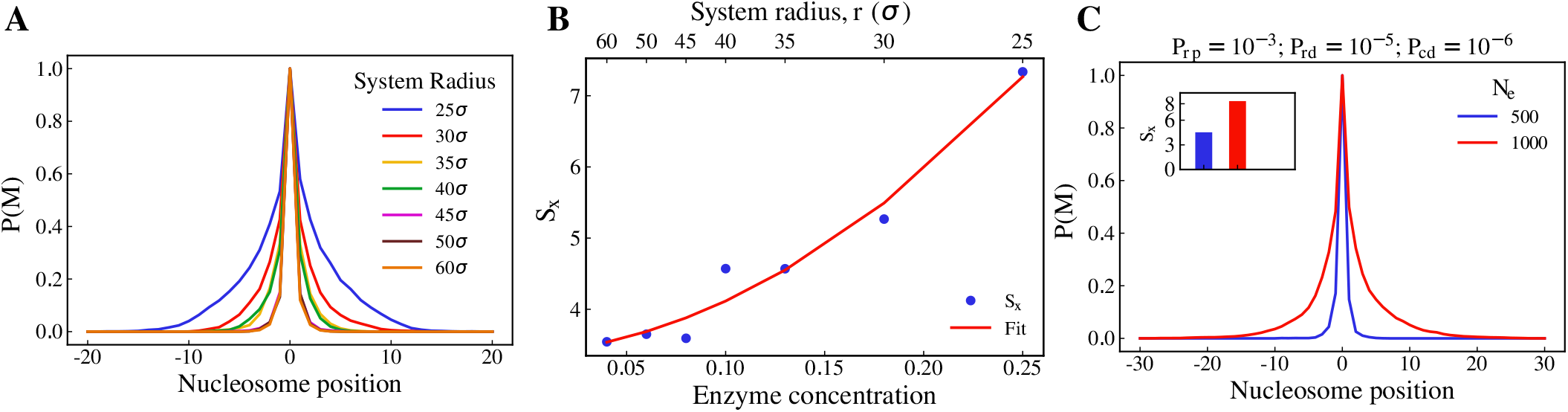
Effect of the total concentration of the constituents on the average methylation profile. The spreading is considerable with an increase in density. (A) The methylation profile is plotted across the lattice positions for systems of various radii (signifying concentration of the system). (B) The standard deviation of the methylation profile(*S*_*x*_) vs total concentration of the system, (C) Comparison of *N*_*e*_ = 500 and *N*_*e*_ = 1000 when *P*_*rp*_ = 10^*−*3^

### C. Comparison of the reaction-diffusion model with a reaction-only model

Models with only kinetics of modification reactions—without diffusion—have been employed to describe the spreading of the modifications. The basic version of such models contains the spreading event of modification from a nucleosome to its neighbor with a rate *k*^+^, and the removal of modification with a rate *k*^*−*^. Such a method was implemented by Hodges *et al*. [52] to predict the modification profile. Similar methods were also used by other groups and the model was extended to incorporate more details into the kinetic events [53, 54]. However, these papers do not consider the effect of diffusion of enzymes.

To understand the effect of diffusion, we compare the results from our reaction-diffusion model with a modified version (see Fig. S6) of the basic reactions-only model [52]. In this modified kinetic model, the nucleation point is always maintained in the methylated state (see section IIB). Fig. 5A shows the methylation profile generated by the Gillespie method for various parameters of 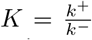, which is the ratio of the rate of the spreading to the rate of removal. The methylation profile widens as we increase *K*. In Fig. 5B, we compare the methylation profiles generated from the diffusion-limited case (*P*_*m*_ = 1) of the reaction-diffusion model to the KMC results. We see that the methylation profile when *P*_*dm*_ = 0.01 matches the methylation profile from the KMC model of *K* = 0.9. However, the methylation profile generated for *P*_*dm*_ = 0.001 is not comparable with either of the two profiles (*K* = 1.3, *K* = 1.5). It suggests that all the profiles generated through the reaction-diffusion model are not reproducible through a reactions-only model, thus illustrating the need to consider diffusion in the model to study this enzyme-mediated spreading phenomenon. To differentiate the profiles generated by KMC from those of reaction-diffusion (RD), we calculated the kurtosis 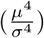 of all the methylation profiles generated. In Fig. 5C, kurtosis values of the methylation profiles are plotted against the standard deviation. We can see that the curve which corresponds to the KMC simulations (No diffusion) stays above the trend of the simulations where the diffusion is accounted (squared red dots). This implies that larger confined domains (high values of standard deviation) with lower kurtosis can be explained only when considering the diffusion of the constituents involved.

**FIG. 5:**
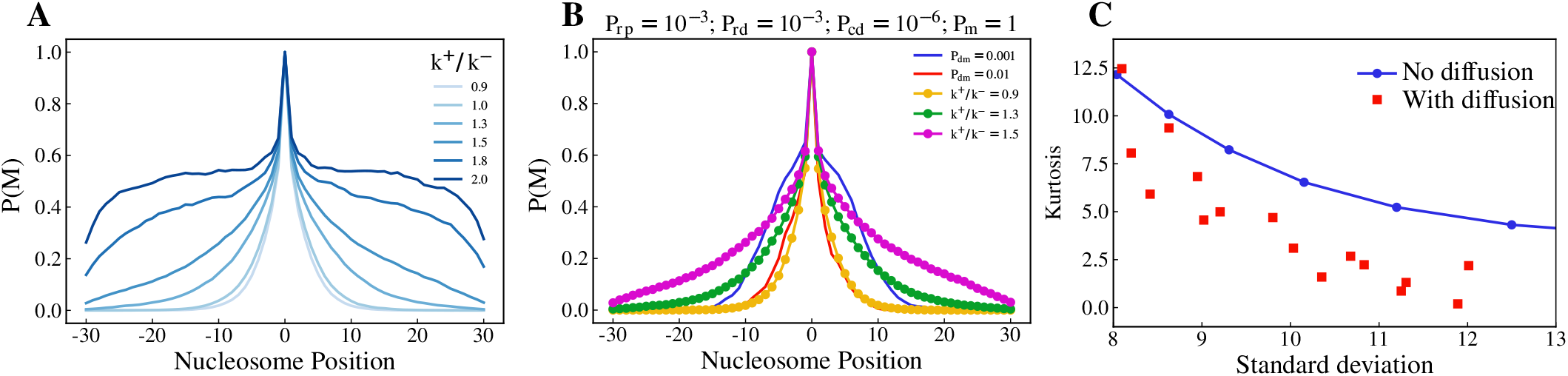
Comparison of methylation profile from the reaction-diffusion particle model to the KMC model. (A) The Methylation profile from the KMC model varied with the parameters. (B) Comparison of the methylation profiles from RD Model with that of KMC. (C) Kurtosis vs Standard deviation of the simulations with and without diffusion.

### D. Confined Methylation profile with a peak can emerge in the absence of a nucleation point

To test the role of the nucleation point, we check a modified version of the reaction-diffusion model (see section II) with two changes: (i) Absence of nucleation point—i.e., the nucleosome at i=0 is like all the other nucleosomes, and it is not constantly in a methylated state, (ii) The enzyme-RNA complex can add methyl marks, with a probability *P*_*w*_, even if the neighbors are not methylated. This is equivalent to an enzyme diffusing and stochastically methylating a random nucleosome, which is in addition to the probability of spreading (*P*_*m*_) to the neighbors by methylated nucleosomes (see Fig. 6A). Fig. 6B shows the plots of methylation profile where kinetic parameters are kept constant, and the curves correspond to different *P*_*cd*_ values ranging from 10^*−*4^ to 10^*−*2^. Similar to the results presented in Fig. 2C, the methylation profile corresponding to the lower *P*_*cd*_ value is wider. Since the nucleosome at *i*=0 is not methylated at all times, the shape of the profile is not the same as that of Fig. 2. However, note that there is a peak emerging at *i*=0. Fig. 6C shows the methylation profiles for different *P*_*w*_ values. Interestingly, as *P*_*w*_ increases, the peak height of the methylation profile increases.

**FIG. 6:**
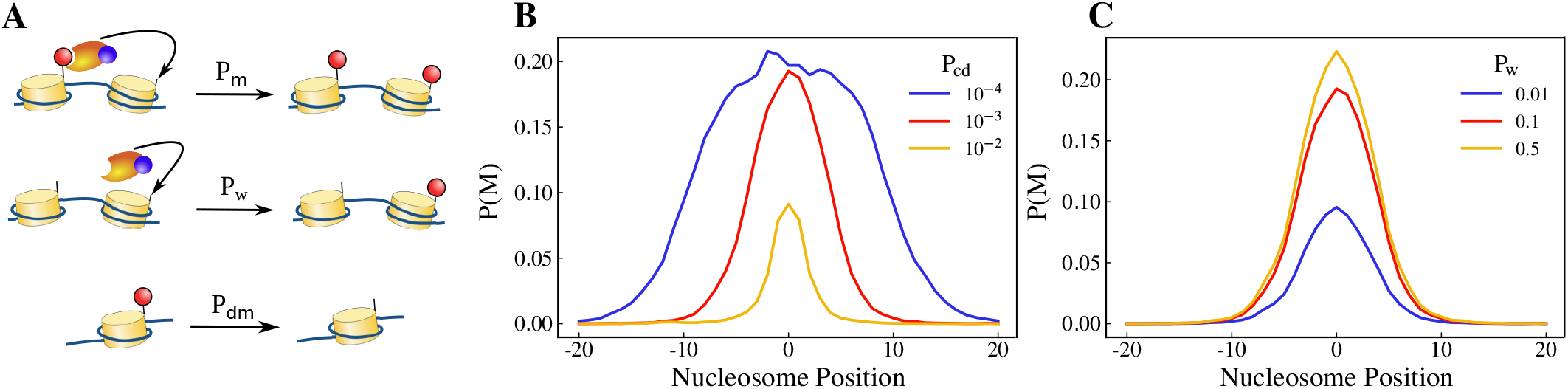
Emergence of nucleation point as a result of the RNA-based mechanism (Model without a nucleation point and a modified probability). (A)Schematic of the nucleosome modification reactions. When Complex particles are in the proximity of unmodified nucleosomes, they can put a mark with probability *P*_*w*_, and if one of its neighbors is modified, this probability will be boosted by *P*_*m*_. (B) Modification profile comparison while varying the rate of complex decay(*P*_*cd*_). (C) Modification profile for different writer probability (*P*_*w*_).

The emergence of the peak at the center of the simulation region without the necessity of a nucleation point can be explained by the RNA particles being produced there. As a result, the enzyme-RNA complex profile peaks around the center, which in turn determines the methylation profile. This suggests that the nucleation points, although meant to be inherited by the daughter cells from the mother cells, can also be a by-product of gene expression from a specific loci. This is achieved by considering the “writer” enzyme property along with “reader-writer” enzymes.

### E. Chromatin polymer folding affects the modification profile and comparison with experimental data

So far, we have assumed the chromatin as a 1-dimensional string. Nevertheless, in cells, chromatin is folded and packed. Each nucleosome will have a few other nucleosomes in proximity (close contact) beyond the two neighbors along the polymer backbone. Some of the recent papers have studied 1-dimensional chromatin but accounted for non-neighboring methylation spreading by modifiying the spreading rates [60, 61, 69]. Some other papers have considered the explicit polymer nature for studying the spread of the modifications [48, 57, 80, 81, 81–83]. Going beyond these models, in this section, we account for chromatin’s folded polymer nature as well as explicit diffusion of constituents and reactions with the nucleosomes. We study the effect of far-away contacts in spreading modifications, and for simplicity, we restrict ourselves to 2D polymer networks. We simulate chromatin as a N-bead self-avoiding walk and random walk polymers (see Supplementary Note). We take an ensemble of configurations from those simulations. For each of these configurations, we simulate the diffusion and reaction of all the constituents (R, E and C) similar to what is done for the linear polymer, and the resulting modification profile is recorded (see section. II). A few such individual methylation profiles from spreading along the self-avoiding walk polymer configurations are shown in Fig. 7D. Note that the individual profiles generated from spreading along the individual configurations do not typically have a highly peaked and symmetrically decaying profile. This implies that we can expect similar profiles whenever the spreading time is less than the polymer relaxation time. Even though the individual profiles do not have a clearly confined domain of modified nucleosomes when we average over an ensemble of configurations, a different profile emerges (see Fig. 7B), in which we can see a confined modification domain. In Fig. 7A, we have compared the profiles generated from a linear lattice with that of polymers with two different degrees of compaction (self-avoiding walk and random walk) while keeping all the other kinetic parameters constant. We can clearly see that the modification spreads more on a random walk polymer than on a self-avoiding polymer. This is expected since each nucleosome in a random walk polymer have many more neighbors. Also, The size of the modification domain decreases as we increase the kinetic parameters *P*_*rd*_ (see Fig. 7B) and *P*_*cd*_ (see Fig. 7C).

**FIG. 7:**
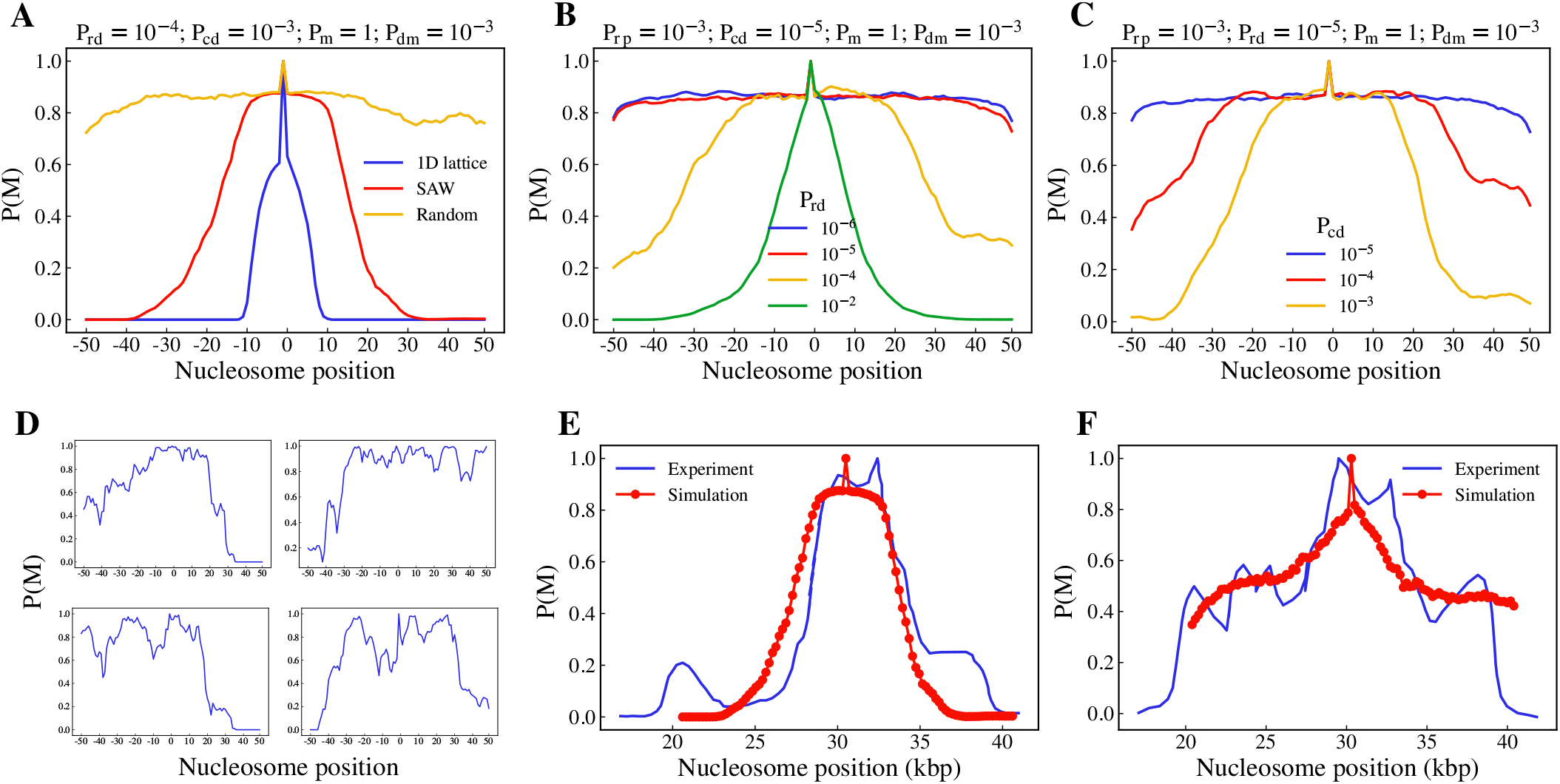
Chromatin compaction engages the spreading. (A)Average Modification profile compared between polymers of different compaction, Average of methylation profile over an ensemble varying with kinetic parameters (B) *P*_*rd*_, (C)*P*_*cd*_, (D) Individual modification profiles from frozen configurations of SAW polymer and random walk polymer, (E) H3K9me3 profile of *hht*2^*G*13*D*^ mutant of the mating-type loci in *S*.*Pombe* compared with the profile generated from an ensmeble of SAW polymers (F) H3K9me3 profile of Wildtype of the above mentioned genome compared with profile generated from random walk polymer

To test whether our simulation results are biologically realistic, we compare them with the experimental data from the Dipiazza*et al*. [75]. They study the spreading mechanism of H3K9me3 from mating type loci (roughly 20kbp) in *S*.*Pombe* using a dominant-negative mutant *hht*2^*G*13*D*^. They establish that the H3K9me3 levels from ChIP-seq are reduced when they use this mutant. We find that the H3K9me3 profile generated from this mutant matches with SAW configurations (see Fig. 7E). Similarly, the H3K9me3 profile of wildtype is comparable with the profile generated from the random walk configurations (see Fig. 7F). This indicates that the profile we obtained in our simulations are realistic.

## IV. DISCUSSION AND CONCLUSION

In this paper, we propose a reaction-diffusion framework to understand the spreading of histone modifications along the nucleosomes. The key difference in our model from other models in the literature is the explicit accounting of the diffusion of proteins/RNAs and mimicking of the RNAi mechanism. We consider a flux of RNA-like (R) particles produced from the nucleation point. These particles diffuse and meet with uniformly distributed diffusing enzyme-like (*E*) particles and then catalyze the propagation of the spreading in our model.

Our main results are as follows: We show that the diffusion of the constituents can play an essential role in deciding the modification pattern. First, we show that the diffusion coefficient and the decay rate of enzyme/protein/RNA constituents set a length scale beyond which the modification cannot spread. This naturally limits the domain size. Second, we show that the distribution of the modification pattern depends on the diffusive aspect, which is different from that of a purely kinetic model. We also simulate how the RNAi mechanism would lead to the emergence of the peak, and we do not have to maintain a nucleation point where a modification-state is consistently maintained with high probability. This suggests that the nucleation sites, which are assumed to be inherited from mother cells, can just be active genes or regions. Some models claim that a nucleation site is required to maintain confined domains, while others suggest a long-range loop-based spreading mechanism mainly to explain the profile [55–58]. It seems more plausible for the switch to be a diffusion-limited mechanism inside the nucleus.

The parameters that control the instantaneous number of these particles are the production and decay probabilities of RNA and the enzyme complex (namely, *P*_*rp*_, *P*_*rd*_, *P*_*cp*_, and *P*_*cd*_). The change in *P*_*rp*_ does not affect the length scale of RNA(see Fig. S7). The modification profile gets wider with the decrease in RNA decay probability (*P*_*rd*_) as well as complex decay probability (*P*_*cd*_). However, this change is not observed in the regime of higher demethylation probability (*P*_*dm*_ = 0.001). Another factor influencing the modification profile in our model is the availability of enzymes. Enzyme limitation has been considered in recent papers by incorporating it into the parameters of kinetic rates. However, enzyme-limitation arises naturally in this model since a given number of enzyme-like particles are part of our model. We varied this enzyme concentration and observed the modification profile getting wider as we increase the concentration. The final aspect we present is the role of chromatin compaction in the spreading of histone marks [48, 57, 80, 81, 81–83]. We show that compaction can help in the spreading process. We compare our model predictions with the H3K9me3 profiles reported in mating-type loci of *S*.*pombe*.

Since most biological processes are determined by an interplay between chromatin organization and reaction-diffusion of proteins, we hope that our model will serve as a starting point in exploring several questions in this direction. The model can be extended to study cooperative (or anti-cooperative) interaction between many different reader/writer enzymes and modifications and the role of many different proteins. Given that many proteins show liquid-like condensation behavior and such condensation is relevant for gene regulation, one has to go beyond the kinetic models and incorporate the reaction-diffusion of proteins into a chromatin model to predict statics and dynamics of the crucial biological mechanisms.

## V. DATA AVAILABILITY

All the data from the work is present in the figures in the manuscript, and supplementary information.

## VI. ACKNOWLEDGEMENT

We acknowledge funding from Science and Engineering Research Board (SERB) (Grant number: CRG/2022/007974). V.M acknowledges the inputs of Sangram Kadam while writing the manuscript.

