## Supplementary Information for "Role of diffusion and reaction of the constituents in spreading of histone modification marks"

### Supplementary Note. Polymer simulation

We considered chromatin as a 2D bead-springs polymer chain of $N$ beads. Each bead in this system corresponds to a nucleosome. We then equilibrated this polymer as a self-avoiding walk and random walk separately. The total energy of the self-avoiding walk polymer is given by,

$$E = \sum_{i=1}^{N-1} E_i^s + \sum_{i,jj>i} E_{LJ}(|r_{ij}|) \tag{1}$$

The beads in this polymer are connected through a harmonic spring, whose potential is given by,

$$E_i^s = \frac{K_s}{2}(|\vec{r_i} - \vec{r}_{i+1}| - r_0)^2 \tag{2}$$

where, $\vec{r_i}$ is the position of the $i^{th}$ beads, $r_0$ is the equilibrium bond length and $K_s$ is the spring constant
All other non-bonded beads interact with LJ potential to achieve volume exclusion.

$$E_{LJ}(r_{ij}) = \begin{cases} 4\epsilon\Big[\Big(\frac{\sigma}{r_{ij}}\Big)^{12} - \Big(\frac{\sigma}{r_{ij}}\Big)^{6}\Big] & r_{ij} < 2^{1/6}\sigma, \\ 0 & r_{ij} \geqslant 2^{1/6}\sigma. \end{cases} \tag{3}$$

where,$\sigma$ is the size of a particle, $\epsilon$ is the strength of the interaction, and $r_{ij}$ is the inter-particle distance.
Then, we performed Brownian dynamics simulations of this model. After the equilibration, we took 100 timeframes such that each frame is $10^5$ timesteps apart from the previous frame. The corresponding position information of the beads were used as frozen configurations for simulating the compaction effects in the reaction-diffusion model. In the case of random walk polymer, the total energy in the system was only the spring potential. Similar procedure was carried out to obtain an ensemble of frozen configurations of random walk polymers.

*Electronic address:
†Electronic address:
‡Electronic address:

**Supplementary Figures**

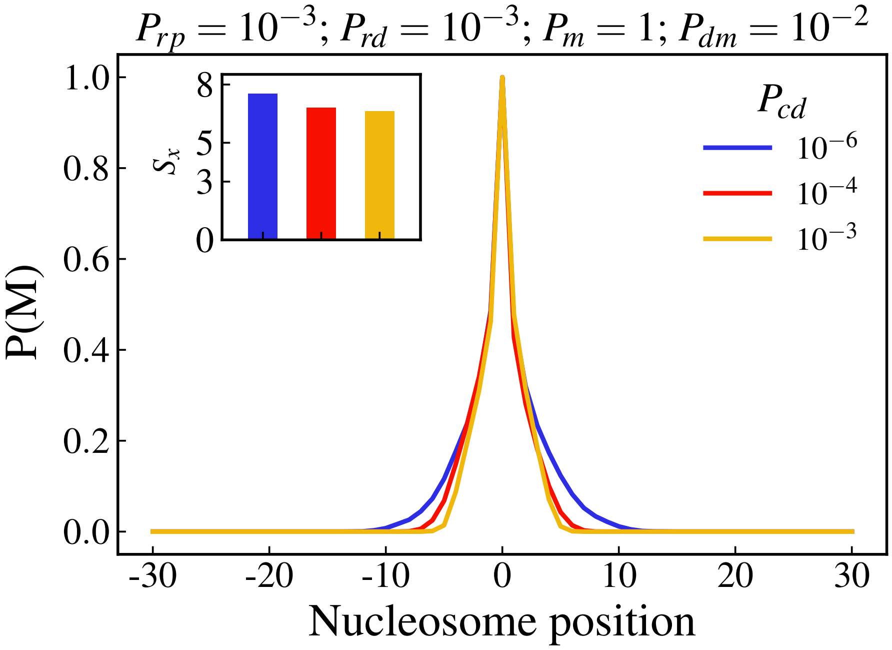


FIG. S1: Effect of $P_{cd}$ in the modification profile, corresponding to the parameters when $P_{dm} = 10^{-2}$. We see no difference between the different profiles, since $P_{dm}$ dominates the effect of complex decay kinetics

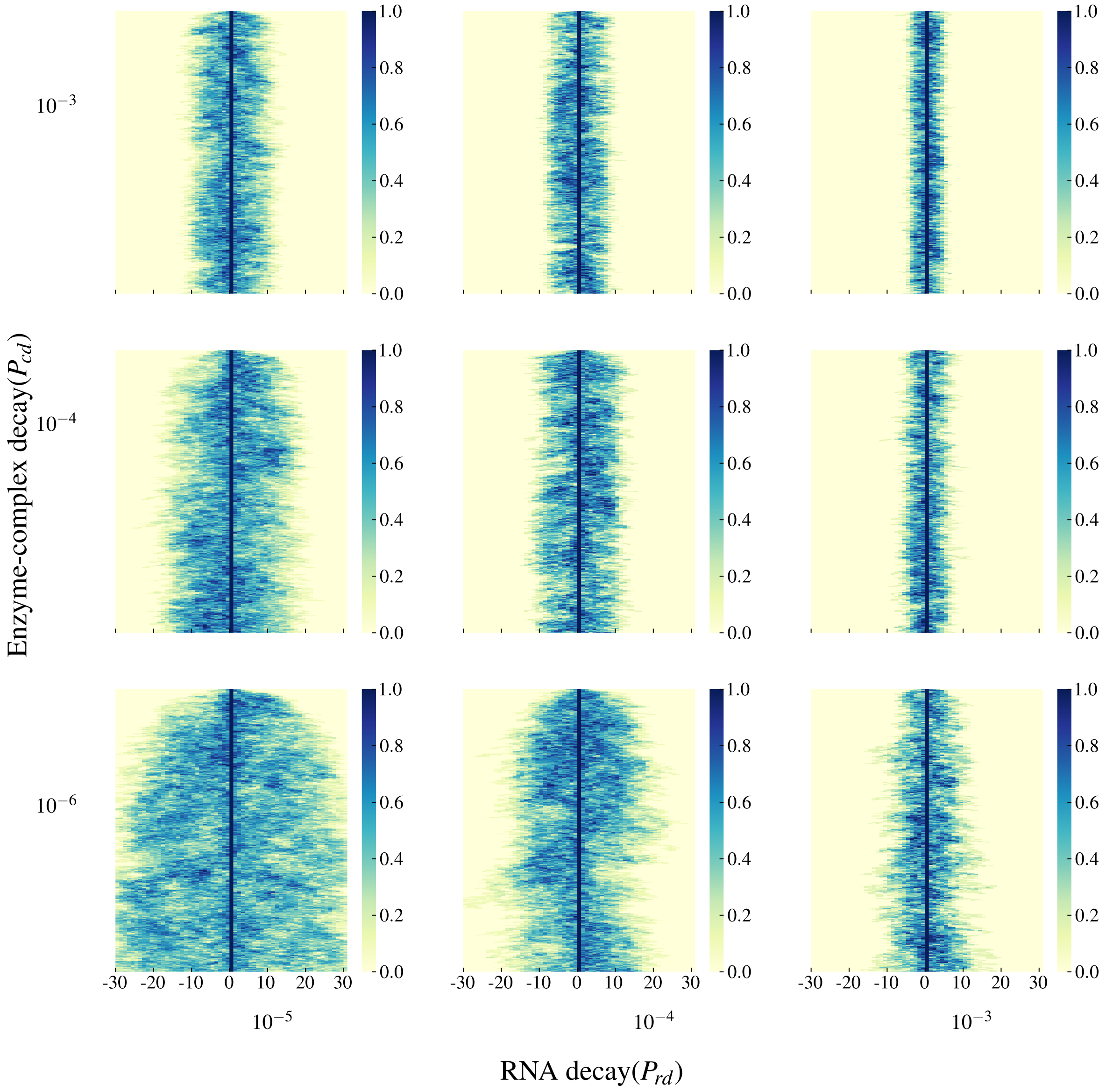


FIG. S2: Phase diagram of time evolution of modification state of nucleosome lattice as we vary the RNA decay parameter ($P_{rd}$) and Enzyme-complex decay parameter ($P_{cd}$). Each individual plot corresponds to a specific $P_{cd}$ and $P_{rd}$ value while all the other parameters are kept constant.

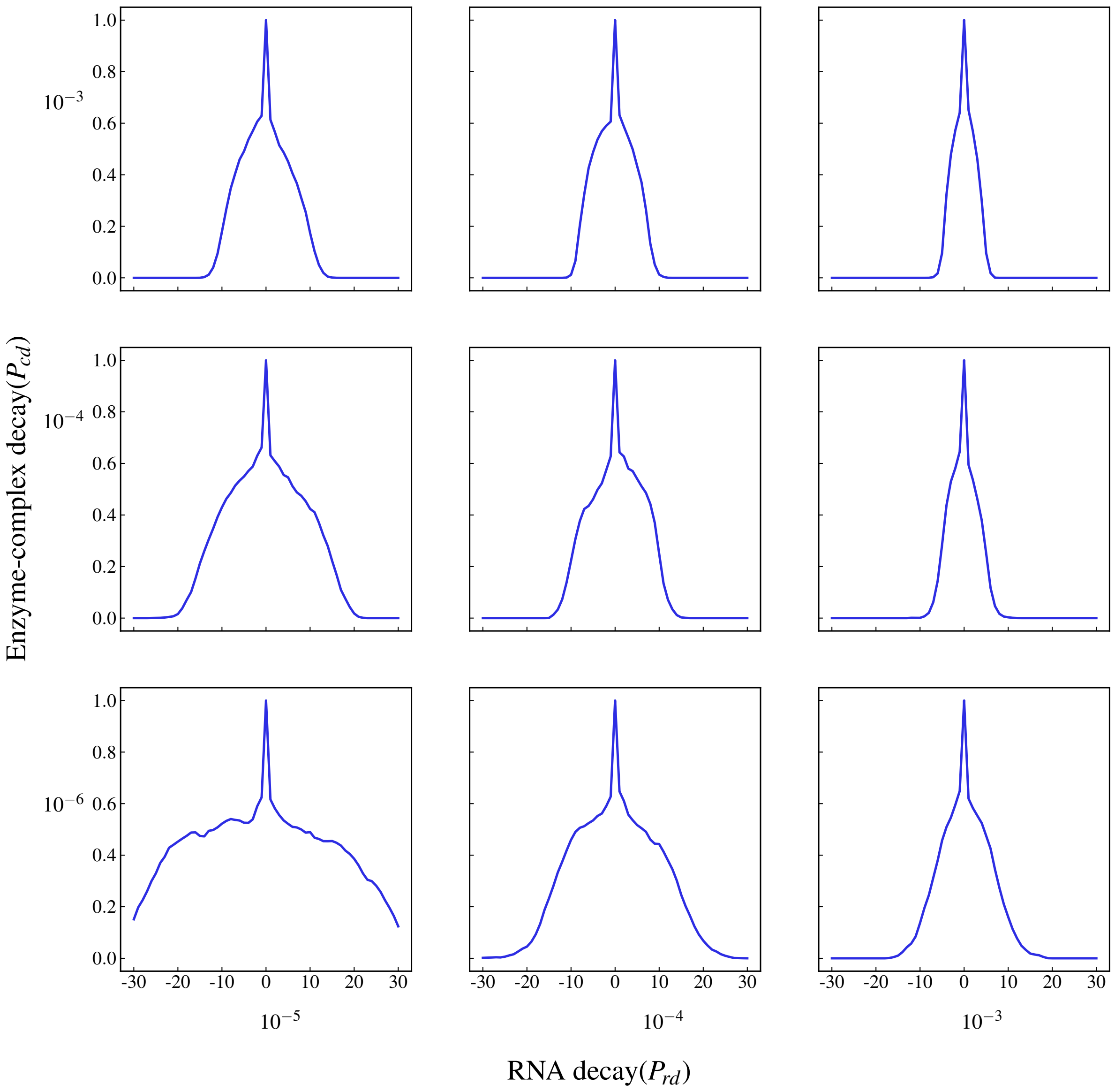


FIG. S3: Phase diagram of modification profile as we vary the RNA decay parameter ($P_{rd}$) and Enzyme-complex decay parameter ($P_{cd}$). Each individual plot shows the steady-state average modification profile corresponding to the parameters mentioned in the axes

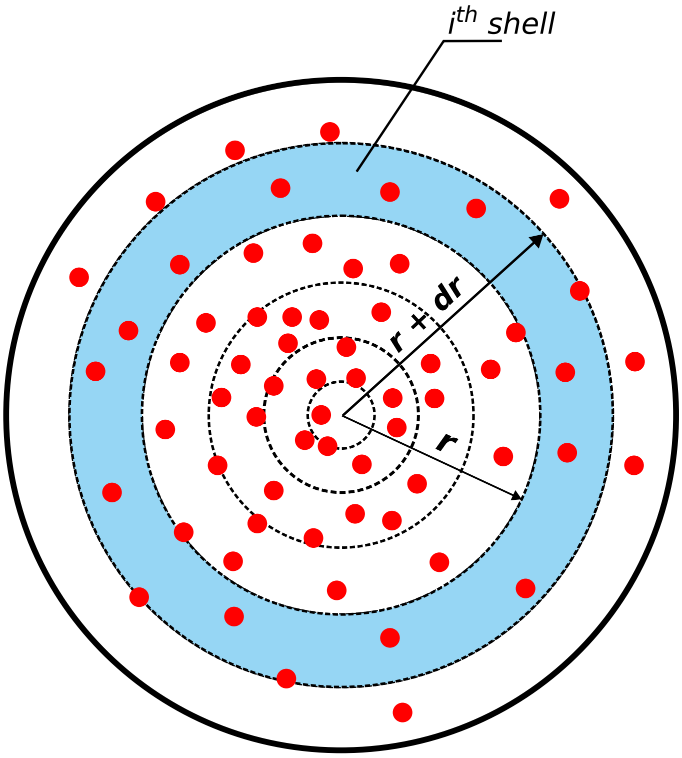


FIG. S4: Schematic of RNA density($\rho_i$) of $i^{th}$ shell. It is the ratio of the number of RNA particles in $i^{th}$ shell (the blue strip of radius $dr$) to the area of the corresponding shell

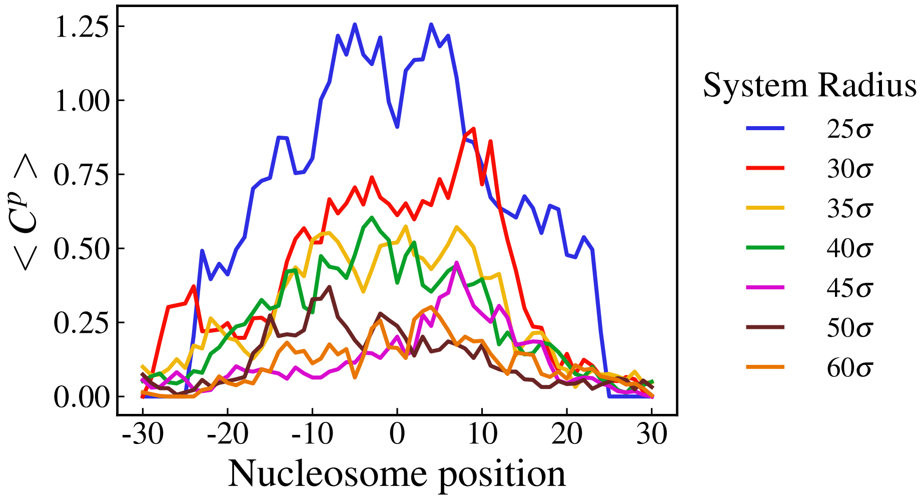


FIG. S5: Average proximity of enzyme-complex($C^p$ plotted against nucleosome position). The different curves correspond to respective system radius. We employed this varied system radius to account for enzyme limitation.

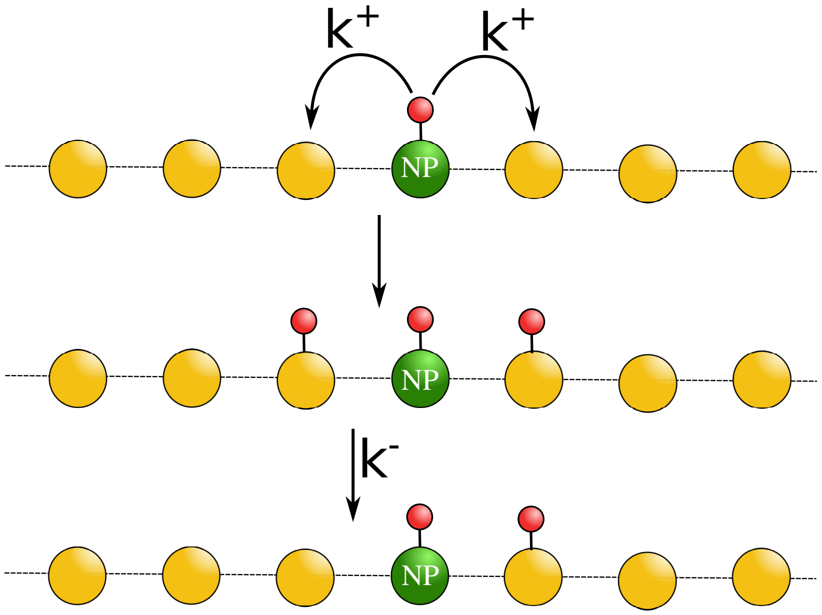


FIG. S6: Schematic of Kinetic Monte-Carlo method (KMC). The nucleation point ($NP$) is kept methylated at all times. The modified nucleosomes can spread their modifications to their neighbours with the rate $K^{+}$ and any modified nucleosome can become unmodified with the rate $K^{-}$

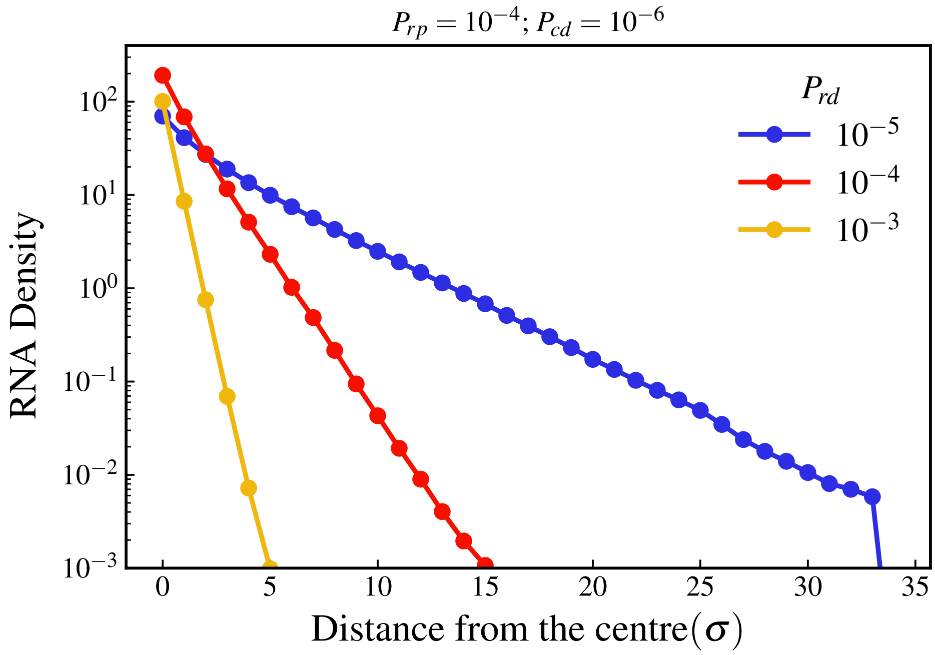


FIG. S7: Spatial distribution of $R$ at the steady state. Variation in number density of R present in the $i^{th}$ concentric shell ($\rho_i$) with $P_{rd}$, 0 starting from the center(see text) when $P_{rp}$= $10^{-4}$. Despite of ten-fold smaller $P_{rp}$, the RNA density remains similar to Fig. 3A

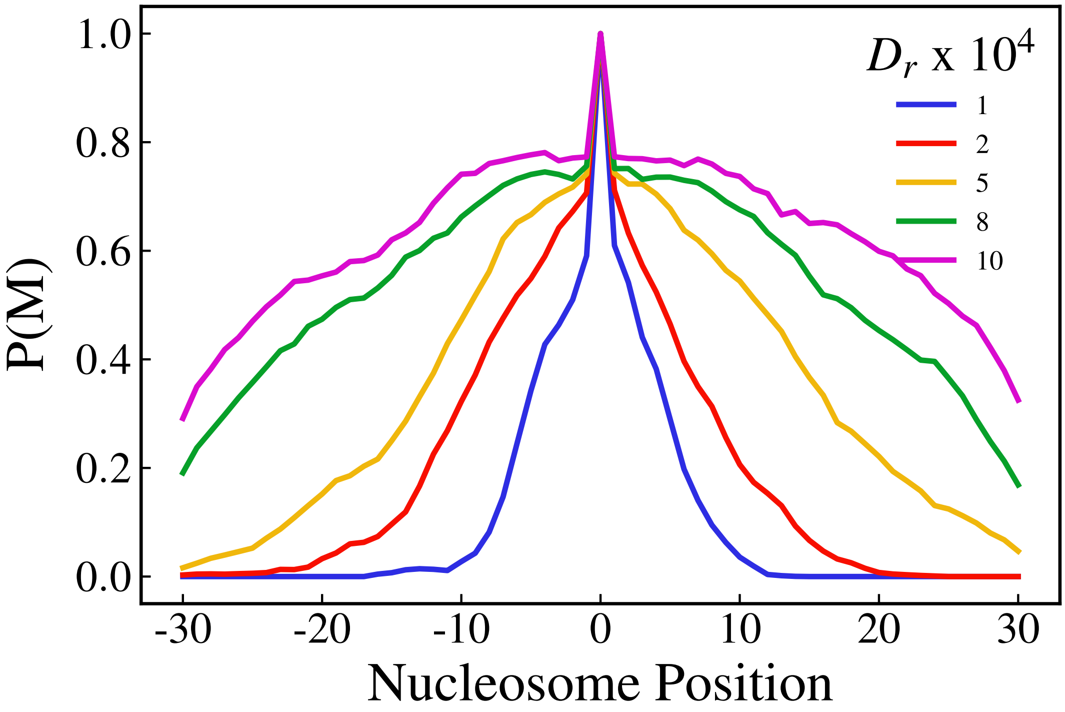


FIG. S8: Methylation profile when we vary the mobility term ($D_r$) in the brownian motion. As we discussed in the main text, the diffusion coefficient and the kinetic parameters together decide the length scale of the modification domains. But we have performed simulations with a constant value of $D_r$ to study the effect of other constituents. However, with a different $D_r$, we would get different sized domains. We have chosen the value $D_r = 2x10^{-4}$, assuming the particle size to be around 5-10 nm, which is biologically relevant.

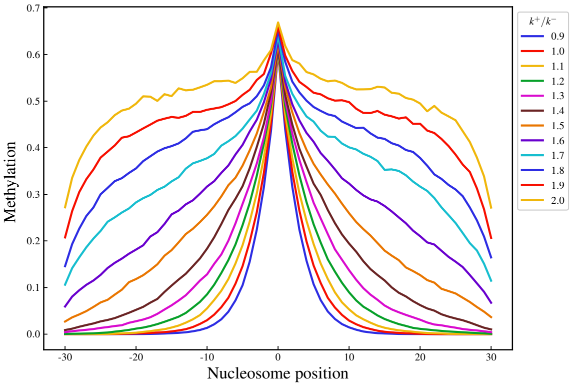


(a)

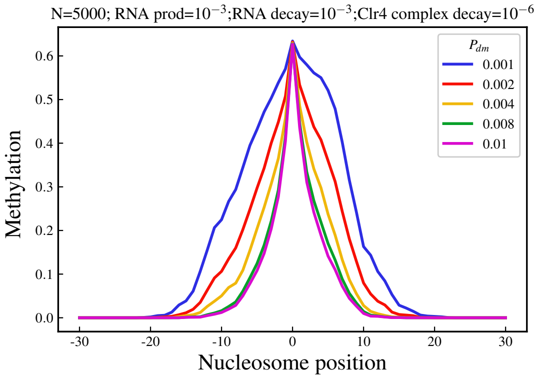


(b)

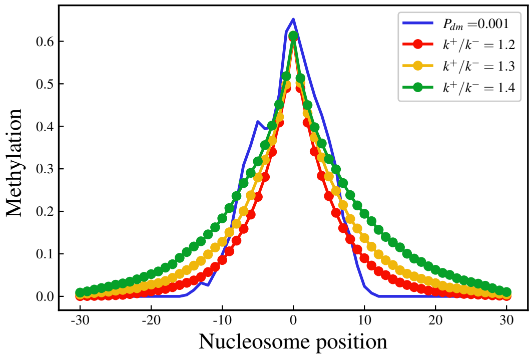


(c)

FIG. S9: The nucleation point is not always maintained at the modified state. This shows that keeping the nucleation point modified throughout the simulation does not qualitatively affect the results. (a) Modification profile of nucleosomes from the KMC model where the conditional nucleation is maintained, (b) Methylation profile from the RD Model where the nucleation point is methylated for 60% time and the different curves represent different probabilities of demethylation($P_{dm}$), (c) Comparison of the methylation profile from $P_{dm} = 0.001$ to different K-values from KMC model to find the equivalent value
